# Minimum requirements for reprogramming and maintaining cell fate in the *Arabidopsis* root

**DOI:** 10.1101/214957

**Authors:** Colleen Drapek, Erin E. Sparks, Peter Marhavy, Tonni G. Andersen, Jessica H. Hennacy, Niko Geldner, Philip N. Benfey

## Abstract

Changes in gene regulation during differentiation are governed by networks of transcription factors. To identify the minimal network for endodermal differentiation in the *Arabidopsis* root, we asked what transcription factors are sufficient to program a non-native cell-type into endodermis. Our results show the transcription factors SHORTROOT and MYB36 have limited ability to reprogram a non-native cell-type (the epidermis) and that this reprogramming is reversible in the absence of additional cues. The stele-derived signaling peptide CIF2 stabilizes SHORTROOT-induced reprogramming. The outcome is a partially impermeable barrier deposited in the sub-epidermal cell layer that has a transcriptional signature similar to endodermis. The trans-differentiation mechanism depends on the expression of genes downstream in the gene regulatory network but is independent of SHORTROOT movement. These results highlight a non cell-autonomous induction mechanism for endodermis that resembles differentiation in many animal systems.

**One Sentence Summary:** SHORTROOT and CIF2 combined can induce and stabilize an endodermis in sub-epidermal cells and do so in a non cell-autonomous manner.

---

Development of multi-cellular organisms requires tight spatiotemporal coordination and stabilization of cell differentiation. The minimal set of components to achieve differentiation is a mystery that has puzzled biologists for more than sixty years (*1*). Transcription factors (TFs) are regulators of gene expression, and are pivotal components of differentiation(*2*). TF-regulated events that underlie differentiation are often simplified in the construction of gene regulatory networks (GRNs)(*3*). The *Arabidopsis* root endodermis has been a useful model for understanding differentiation and has a defined GRN, as well as signature features of differentiation(*4*). Differentiated endodermis has lignified cell-wall impregnations called Casparian strips as well as developmentally delayed secondary cell-wall depositions of suberin lamellae(*5*, *6*). Both of these features are thought to contribute to the endodermis’ physiological role acting as a barrier to the external environment (*5*, *6*).

The key TFs of the endodermis GRN are SHORTROOT (SHR), SCARECROW (SCR) and MYB36. These TFs form a transcriptional cascade with SHR as the “master regulator” activating SCR and subsequently MYB36(*4*). SHR is expressed in the root vasculature (stele) and moves into the neighboring cells where it initiates the asymmetric cell division that separates the endodermis from the cortex (collectively the ground tissue)(*7*). Both the *scr* and *shr* mutants contain only one layer of ground tissue, with *shr* being devoid of lignin deposition and *scr* having a sporadic lignification pattern(*8*, *9*). MYB36, which is downstream of SHR and SCR, regulates expression of CASPARIAN STRIP ASSOCIATED PROTEINS (CASPs) (*10*–*12*). Mutations in MYB36 result in delayed apoplastic barrier formation (*10*, *11*). Proper endodermis formation also relies on the receptor-like kinase SCHENGEN3 (SGN3), which is downstream of SHR(*13*, *14*). SGN3 seals the Casparian strip upon activation by the stele-derived ligands CASPARIAN STRIP INTEGRITY FACTOR 1 and 2 (CIF1/2) (*15*, *16*).

Ectopic expression of *SHR* or *MYB36* results in few cells with ectopic lignin deposition (*11*, *17*). It remains unclear what is the minimum pathway required for endodermal differentiation that can instruct a non-native cell type to develop endodermal features. To determine if additional components are required, we individually expressed *SHR*, *SCR*, and *MYB36* in the epidermis using the *WEREWOLF (WER)* promoter (*18*). We used expression of CASPs, and the deposition of lignin and suberin as markers for endodermal differentiation. Consistent with previous results, we found that epidermal-expressed SHR induces stochastic expression of lignin in a few epidermal cells (Fig. 1A-B). We further found that CASP1 (Fig. 1C-D) and suberin (Fig. S1A-B) are in a small subset of epidermal cells and, occasionally, in the supernumerary ground tissue layers that are present in these lines as a result of the overexpression of *SHR* (*17*). Epidermal expression of *SCR* was unable to induce CASP1, lignin or suberin (Fig. S1C-F). As was previously reported for lines driving *MYB36* under a ubiquitous promoter (*11*), we found epidermal-expressed *MYB36* also had limited ectopic CASP1 and lignin deposition in the epidermis (Fig. 1E-F), as well as suberin deposition (Fig. S1E,G). The ectopic presence of suberin in these lines was unexpected as the *myb36* mutant has an over-abundance of suberin (*11*). We conclude that *SHR* and *MYB36*, but not *SCR*, are able to stochastically generate endodermal features in the epidermis.

**Fig. 1.**
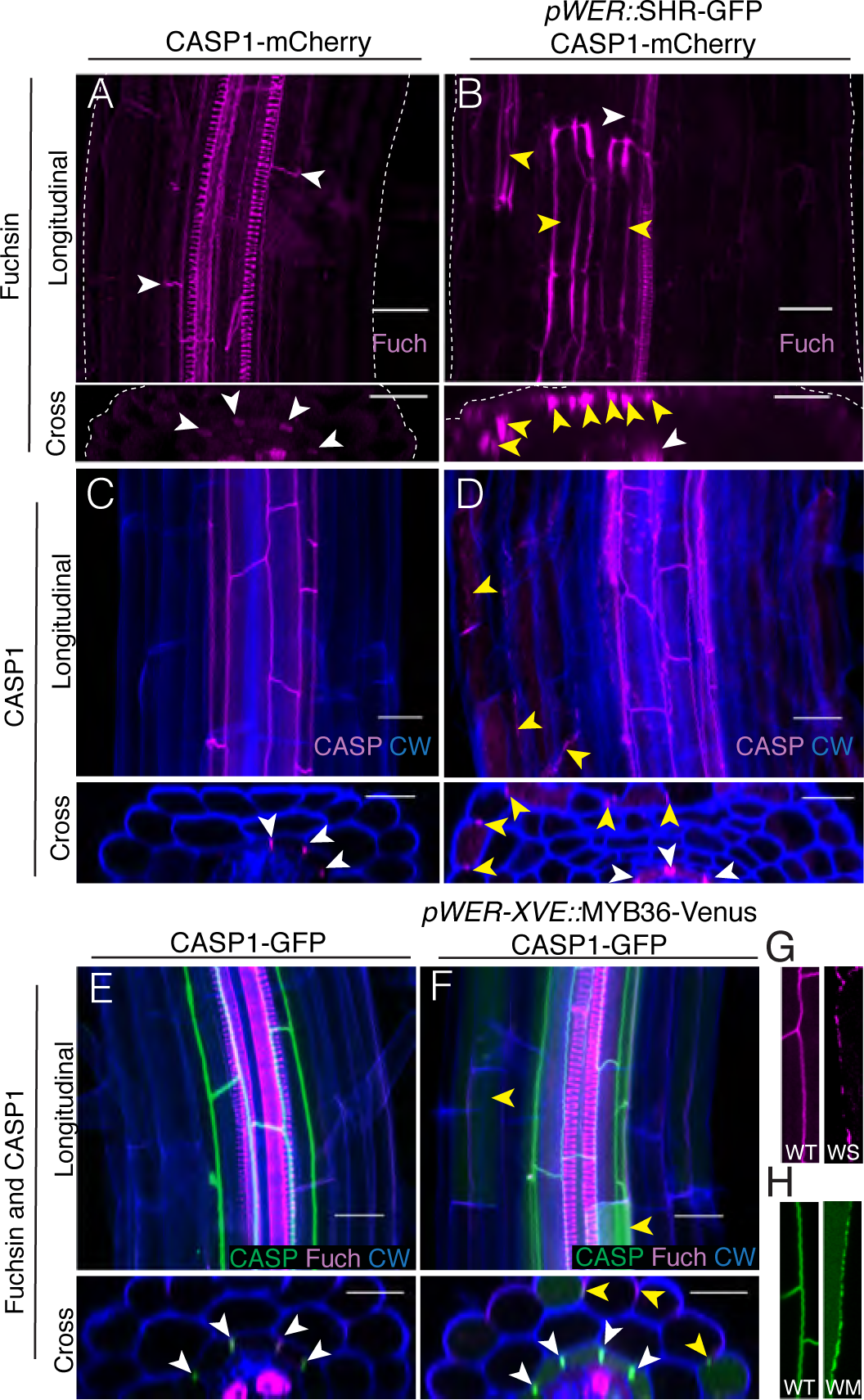
*SHR* and *MYB36* can induce ectopic endodermal features in the epidermis. (A) Max projections of longitudinal and cross section views of CASP1-mcherry (wild-type) and (B) *pWER::*SHR:GFP x CASP1-mCherry Basic Fuchsin (Fuch) staining. White arrowheads indicate endodermal lignin, yellow arrowheads indicate ectopic lignin. Dashed lines in (A-B) indicate outline of the root. Note: fixing method in (A-B) eliminates fluorophore signal. (C) Sum projections of longitudinal and cross section views of cleared CASP1-mCherry (wild-type) and (D) *pWER::*SHR:GFP x CASP1-mCherry roots stained with calcofluor white (CW). White arrowheads indicate endodermal CASP1, yellow arrowheads indicate ectopic CASP1. (E) Sum projections of longitudinal and cross section views of cleared CASP1-GFP (wild-type) and (F) *pWER-XVE::*MYB36-Venus x CASP1-GFP roots germinated on 10µM β-estradiol stained with fuchsin (Fuch) and calcofluor white (CW). White arrowheads indicate endodermal lignin/CASP1, yellow arrowheads indicate ectopic lignin/CASP1. (G) Zoomed in sum projections of endogenous Casparian strip domain in CASP1-mCherry in wild-type (WT) or ectopic Casparian strip domain in *pWER::*SHR:GFP (WS). (H) Zoomed in sum projections of endogenous Casparian strip domain in CASP1-GFP in wild-type (WT) or ectopic Casparian strip domain in *pWER-XVE::*MYB36-Venus x CASP1-GFP (WM). Scale bars = 20µM.

The ectopic Casparian strip-like structures in the epidermis do not resemble the native Casparian strip. Instead, they have a “beads on a string” phenotype similar to the *sgn3* receptor mutant (Fig. 1 G-H) (*13*). It was previously reported that *SGN3* expression is significantly down-regulated in *shr* but is relatively unaffected in *scr* or *myb36* (*10*, *14*). Recently, the peptide CIF2 was identified as a ligand for SGN3. CIF2 is produced in the stele and its expression is independent of SHR (*14*). We hypothesized that the “beads on a string” phenotype of the ectopic Casparian strip-like structures was a result of present but inactive SGN3 as CIF2 would not be able to reach the epidermal cell layer from the stele. To test this hypothesis, we treated epidermal-expressed *SHR* seedlings with 100 nM CIF2. This treatment resolved the “beads on a string” phenotype, however, the addition of CIF2 also induced polarized CASP1 in 80–100% of the cells in the sub-epidermal layer (Fig. 2A-B). These cells also deposit lignin (Fig. 2C-D) and suberin (Fig. S2A-B). We confirmed that cells ectopically expressing *SHR* can induce *SGN3* (Fig. S3A-C) and that this fate change is not due to *SGN3* alone, since epidermal-expressed *SGN3* in the *sgn3-3* background was not capable of reprogramming the epidermis with or without CIF2 (Fig. S3C-F). Ectopic Casparian Strip-like structures in epidermal-expressed *MYB36* roots also have a “beads on a string” phenotype (Fig 1H). However, since *SGN3* expression is unaffected in a *myb36* mutant (*10*)(Fig. S3G-H), we hypothesized that expression of *MYB3*6 is not sufficient to activate *SGN3*, and thus CIF2 treatment would not seal ectopic Casparian strip-like domains. Consistent with our hypothesis, treatment with CIF2 did not seal ectopic domains nor did it induce significantly more epidermal cells to form Casparian strips (Fig. S4A-E). Plants with epidermal-expressed *SCR* show no markers of trans-differentiation even when treated with CIF2 (Fig. S4F-I). Although neither epidermal-expressed MYB36 nor SGN3 is sufficient to induce reprogramming in the presence of CIF2, they are both required for the reprogramming phenotype observed in epidermal-expressed SHR plants (Fig. S4 J-O).

**Fig. 2.**
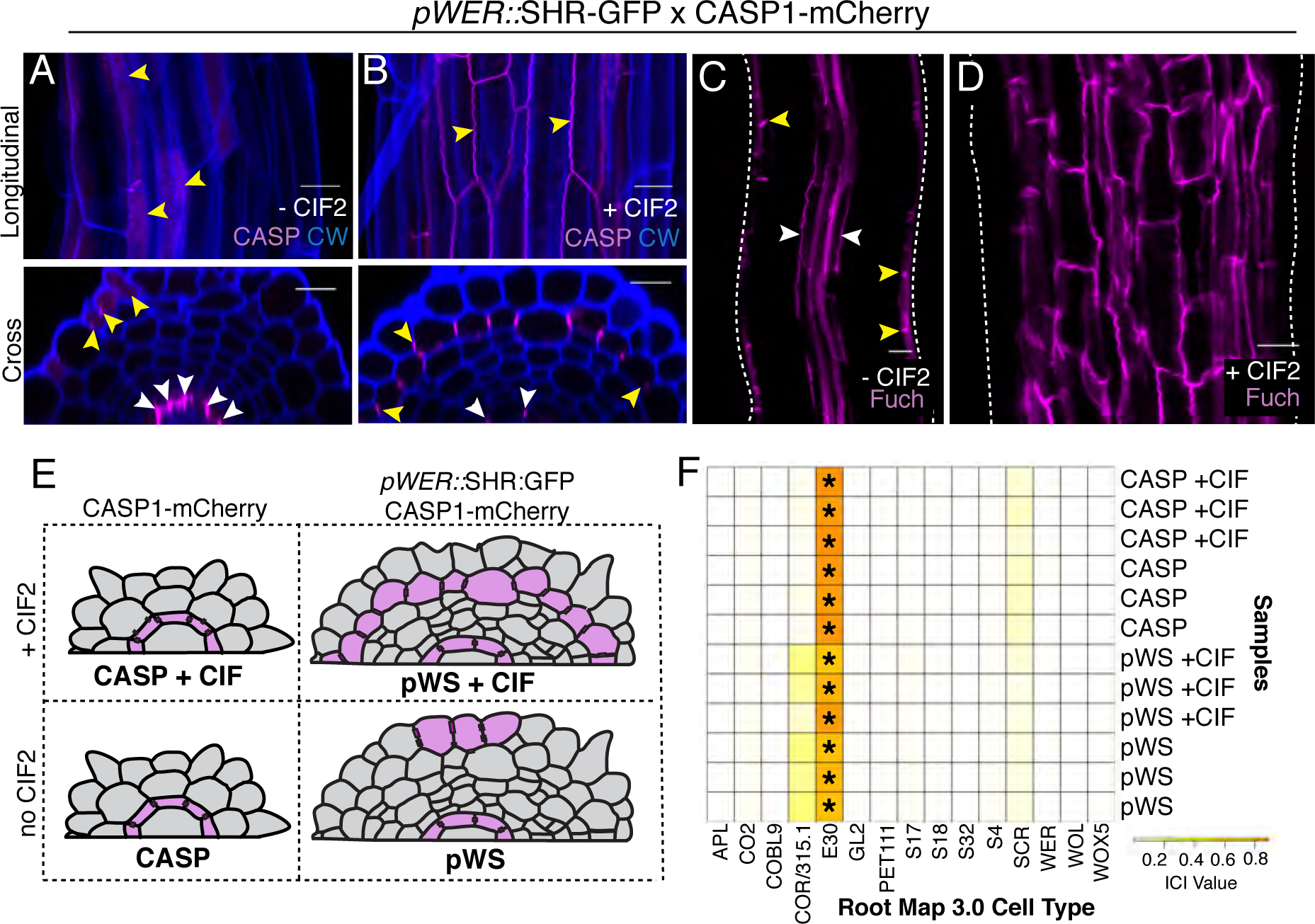
CIF2 induces reprogramming of the sub-epidermal layer. (A) Sum projection of longitudinal and cross section views of the epidermis and ground tissue layers of cleared *pWER::*SHR:GFP x CASP1-mCherry roots without CIF2 and (B) with 100 nM CIF2. Yellow arrowheads in (A-B) indicate examples of ectopic CASP1. White arrowheads in (A-B) indicate native CASP1. CW=calcofluor white. (C) Sum projection of Basic Fuchsin (Fuch) stained *pWER::*SHR:GFP x CASP1-mCherry roots without CIF2 and (D) with 100 nM CIF2 (sum projection of the sub-epidermis). White dashed line indicates root outline. Scale bars = 20µM. (E) Schematic of four samples in RNAseq experiment. (F) Index of cell identity (ICI) for three replicates of samples in (E) across RootMap 3.0. Shade indicates ICI value. *= ICI p-val<0.001.

We then asked if the sub-epidermal cell layer resembles an endodermis or simply produces a Casparian-strip like structure in a cortex cell. We performed RNAseq on FACS-sorted mCherry-positive cells. CASP1-positive cells were analyzed from wild-type plants expressing CASP1-mCherry and from plants with epidermal-expressed *SHR* (*pWER::*SHR:GFP;CASP1-mCherry) with or without CIF2 treatment (Fig. 2E). Note, that in all cases, cells from the endogenous endodermis express CASP1 and thus are analyzed along with the ectopically expressing cells. However, the number of ectopically expressing cells differs dramatically with and without CIF2 in epidermal-expressing SHR lines. Interestingly, treatment with CIF2 yielded only two genes differentially expressed compared to the untreated controls in both genotypes (Table S1). This suggests that CIF2 acts primarily in a post-transcriptional manner on CASP1-positive cells. In a comparison of wild-type and epidermal-expressed *SHR*, there were 172 differentially expressed genes with both receiving CIF2 treatment and 62 differentially expressed genes without treatment (Fig. S5 A, Table S1). Of the 172 differentially expressed genes, only 12 were down-regulated in epidermal-expressed *SHR* as compared to wild-type suggesting only a few endodermis-associated genes are missing from the reprogrammed layer (Table S1). Many of the genes up-regulated in the epidermal-expressed *SHR* lines show enriched expression in the native cortex (Fig. S5B), indicating that this approach has captured reprogrammed cells that retain some of their former cell state. To determine the identity of the trans-differentiated cells, we carried out a Principal component analysis comparing our obtained profiles to those from high-resolution RNAseq cell type-specific profiles (RootMap 3.0) (Fig. S5A) (*19*). The epidermal expressed *SHR* with CIF2 treatment clustered closely with wild-type CASP1-positive cells (no CIF2) and with the endodermis marker E30 as compared to the cortex marker COR (315.1). This suggests that the reprogrammed cells most closely resemble mature endodermal cells even though they retain a minor amount of cortex gene expression.

We also calculated an Index of Cell Identity (ICI) to characterize cell identity based on the weighted expression of marker genes defining that tissue (*20*). We calculated Spec scores for marker genes using the RootMap 3.0 data, and calculated the ICI for CASP1-positive (wild-type) and epidermal-expressed *SHR* samples, both with and without CIF2 treatment. We found that the reprogrammed cells significantly associate with endodermal identity (E30), but retain a low, non-significant, level of cortex identity (COR/315.1) (Fig. 2F). These analyses indicate that the epidermal-expression of *SHR* is sufficient to reprogram to an endodermal cell fate and predominantly turn off the native cortex gene expression. Interestingly, cell fate was effectively switched in CASP1-positive cells regardless of CIF2 treatment, suggesting that CIF2 functions post-transcriptionally to stabilize cell fate.

The sporadic nature of reprogrammed epidermal cells prior to CIF2 treatment suggested that expression of the downstream targets was unstable. To analyze the dynamics of downstream targets, we live-imaged CASP1 in plants with epidermal-expressed SHR for 24 hours. We found that the ectopic CASP1 intensity fluctuates over time (Movie 1, Fig. 3A-B). Interestingly, the CASP1-positive cells were not necessarily the same cells assembling lignin (Fig. 3C). This could indicate that reprogramming is not a linear process in which the steps follow a predetermined order, or could be a result of the instability of CASP1. Consistent with the stochastic expression of CASP1, *SCR* is also expressed in some but not all epidermal meristematic cells in epidermal-expressed *SHR* roots (Fig. S5 A-C). By contrast, *MYB36* expression was occasionally detectable in the epidermis, but primarily induced in the sub-epidermal cell layer (Fig S5 D-F), suggesting that *SHR* may have a non-cell-autonomous role in regulating *MYB36* expression.

**Fig. 3.**
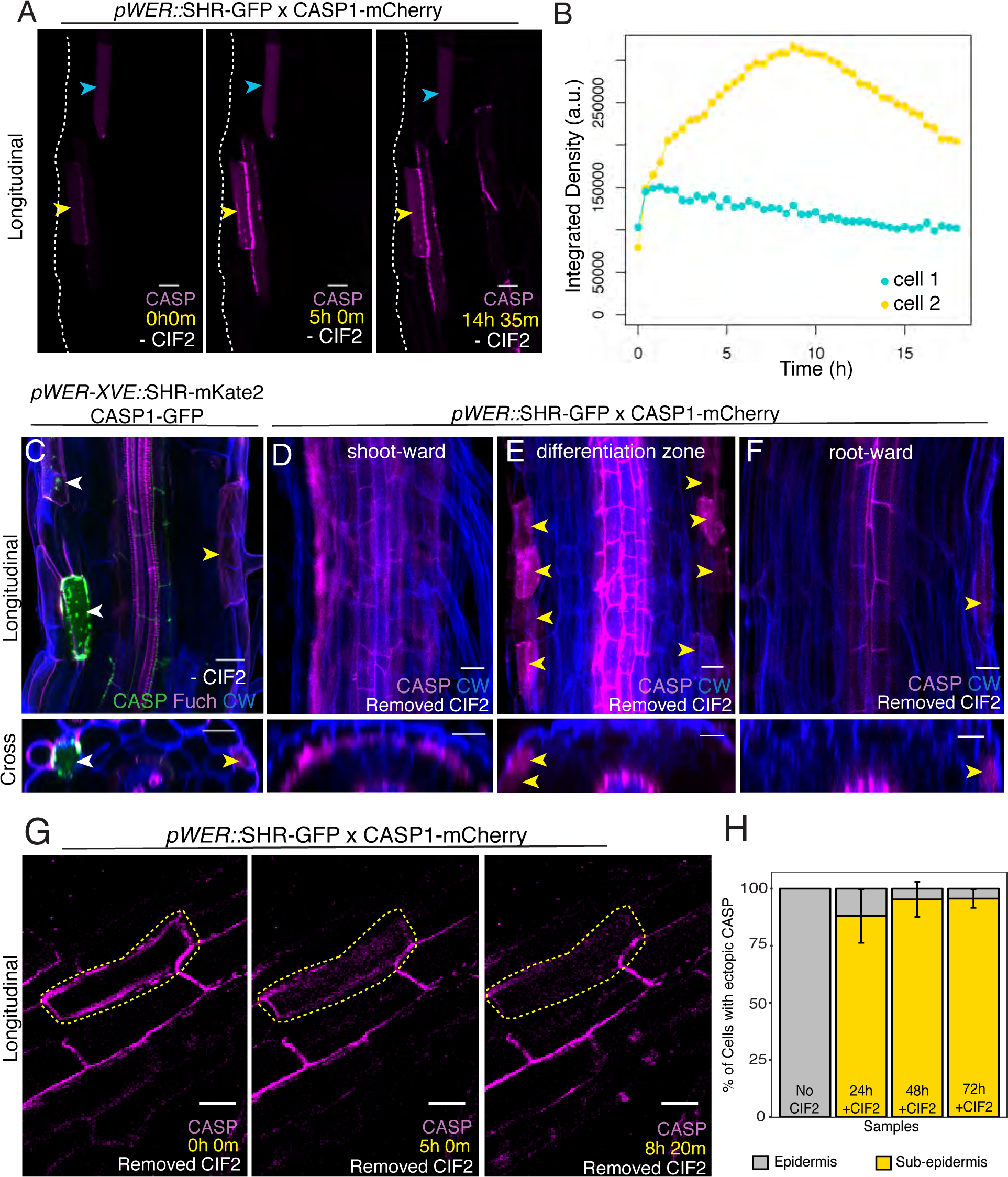
Reprogramming is unstable and non-linear. (A) Max projection frames from live imaging of *pWER::*SHR:GFP x CASP1-mCherry. Cyan and yellow arrowheads indicate cells with fluctuating CASP1 intensity. (B) Changes in intensity over time of cell 1, cyan arrowhead in (A), and cell 2, yellow arrowhead in (A). (C) Max projection of longitudinal and cross section views of *pWER-XVE::*SHR-mKate2 x CASP1-GFP germinated on 10µM β-estradiol cleared and stained with calcofluor white (CW) and Basic Fuchsin (Fuch). White arrowheads indicate cells with both ectopic CASP1 and lignin. Yellow arrowheads indicate cells with ectopic lignin but not CASP1. (D-F) Sum projection of a cleared and stained *pWER::*SHR:GFP x CASP1-mCherry roots germinated on and then removed from CIF2 for 24h. CW= calcofluor white.(F) Shoot-ward (most mature) section, (E) section within differentiation zone prior to CIF2 removal and (F) section containing dividing/elongating cells prior to CIF2 removal. Yellow arrowheads indicate ectopic CASP1 in epidermis. (G) Sum projection frames from live imaging of *pWER::*SHR:GFP x CASP1-mCherry roots germinated on and then removed from CIF2. Yellow dashed outline follows cell that loses polarization and intensity of CASP1. (H) Quantification of the percentage of cells expressing ectopic CASP1 in the epidermis layer or sub-epidermis layer in random sections of roots treated with CIF2 for 24–72h. Standard t-test P-val of treatments vs. no CIF2 are 24h=0.15, 48h=0.029 and 72h=0.013. N=5. Error bars represent standard deviation. Scale bars = 20µM.

To determine if CIF2-induced expression and localization of ectopic CASP1 resulted in commitment to endodermal cell fate, we transferred five-day-old seedlings germinated in the presence of CIF2 to plates without CIF2 for 24h. Under these conditions, the most mature (shootward) cells retained their CASP1 expression (Fig. 3D). However, in a small region of the differentiated tissue closer to meristematic tissues, CASP1 became diffuse, depolarized and abundant in the epidermis (Fig. 3E). Rootward of this region, which contains cells that were dividing prior to transfer, the root appears similar to one that is untreated with CIF2 (Fig. 3F). In order to understand the dynamics of CASP1 in the unstable region (Fig. 3E), we live imaged differentiated sub-epidermal cells of plants removed from CIF2 treatment. We saw a loss of intensity and polarity of established CASP1 in the sub-epidermal cells within this region (Fig. 3G, Movie 2). Finally, we asked if the fate of the few cells expressing CASP1 in the epidermis layer was stable after CIF2 treatment. We found significantly fewer cells expressing CASP1 in the epidermis after 48h and 72h of CIF2 treatment (Fig. 3H). Taken together, these results suggest that cells no longer dividing or elongating have a narrow window in which they can reverse CASP1 expression and localization. This is similar to fate reversal in induced-pluripotent stem cells and provides evidence that expression of differentiation markers is reversible *in planta* (*2*).

The epidermal-expression of *SHR* induces cortex-like supernumerary layers due to the movement of SHR within the epidermal stem cells(*17*, *21*). Since CIF2-induced trans-differentiation occurs in the sub-epidermal layer, this raised the possibility that SHR movement is required for trans-differentiation. To test this hypothesis, we expressed SHR in the epidermis with a nuclear localization tag to sequester it to the nucleus and inhibit movement (Fig 4A-F). Consistent with our results with the non-nuclear-localized SHR, the nuclear-localized SHR lines display only a few cells in the epidermis that express or localize CASP1, lignin or suberin (Fig. 4A-B, Fig. S7A). Upon CIF2 treatment, 70–90% of the cells in the layer that would normally be cortex contain CASP1, lignin and suberin (Fig. 4 D-F, Fig. S7B), indicating trans-differentiation does not require SHR movement.

**Fig. 4.**
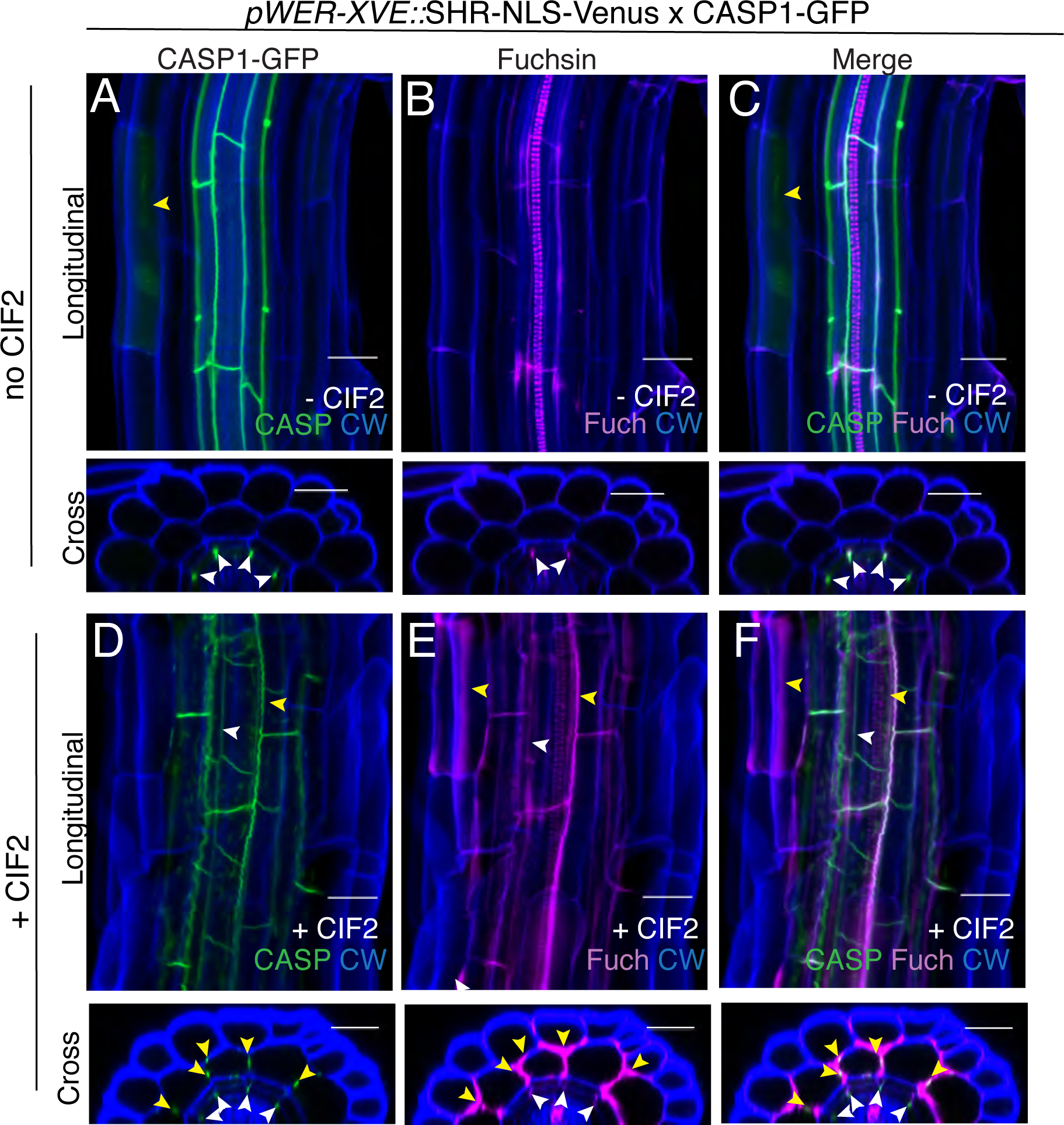
Reprogramming does not require SHR movement. (A) Sum projections of longitudinal and cross section views of cleared *pWER::*XVE:SHR-NLS-Venus x CASP1-GFP roots germinated on 10µM β-estradiol without CIF2, (B) Basic fuchsin stained (Fuch), merge (C). (D) Sum projection of longitudinal and cross sections views of cleared pWER::XVE:SHR-NLS-Venus x CASP1-GFP roots germinated on 10µM β-estradiol and 100nM CIF2, (E) fuchsin stained (Fuch), merge (F). White arrowheads indicate endogenous CASP1/lignin. Yellow arrowheads indicate ectopic CASP1/lignin. CW = calcofluor white. Scale bars = 20µM.

Epidermis-expressed SHR combined with CIF2 result in Casparian strip-like features, suggesting that this structure might form an impermeable barrier. We tested the effectiveness of the trans-differentiated layer to act as a symplastic barrier by measuring FDA penetration (*22*) and as an apoplastic barrier by propidium iodide penetration (*23*). The sub-epidermal layer in both p*WER*::SHR-GFP alone and p*WER*::SHR-GFP with CIF2 had similar FDA penetration rates, suggesting that the suberin deposited in this layer is not a complete symplastic barrier (Movie S1, S2). Propidium iodide penetration was partially inhibited in the sub-epidermal cell-layer (Fig. S8A-B), suggesting that the ectopic endodermal cell wall components form a limited barrier that is not as effective as native endodermis.

Our results establish a previously unappreciated role for receptor-like kinase signaling in stabilizing transcription networks underlying differentiation, a process that was assumed to function far downstream of the events controlled by the early “master regulator” *SHR*. How *SHR* and CIF2 can overcome native gene expression to reprogram the differentiation pathway remains a mystery. Interestingly, the formation of root hairs, a hallmark of differentiation in epidermal cells, was never observed when ectopic CASP1 was present. This may be due to expression conferred by the *WER* promoter, which is active in all meristematic epidermal cells but only in non-hair cells in differentiated tissue (*18*). Alternatively, this may be evidence for differentiation being a mutually-exclusive process. In lines expressing SHR under the *COBL9* promoter, which is active in mature hair cells (*24*), no ectopic CASP1 was observed. This suggests *SHR* must be expressed prior to the onset of epidermal differentiation to facilitate reprogramming (Fig. S10 A-B). Epidermal-*SHR* can also induce ectopic *MYB36* in sub-epidermal cells (Fig. S6D-F) and *SHR* reprogramming is independent of its movement (Fig. 4D-F). The most parsimonious explanation for these results is that there is a mobile cue or factor downstream of *SHR* that activates *MYB36*, implying another stele-derived factor in endodermal differentiation. This is supported by a recent publication demonstrating that a stele-derived cue other than *SHR* can activate expression of the ground-tissue associated enhancer trap J0571(*25*). Together, these data support a model in which the primary differentiation cues for the endodermis emerge from the neighboring tissue, the stele. There are several examples of neighboring cells providing differentiation cues in animals, including the classic cases of TGF-ß and FGF signaling in endodermal-induction of mesoderm tissue in vertebrates as well as EGF signaling in drosophila eye development (*26*, *27*). Finally, we note the similarity of this reprogrammed layer to the exodermis, a waterproof sub-epidermal layer present in most vascular plants, but lacking in *Arabidopsis* (*28*). We can only speculate that homologs of these components may play a role in exodermis formation in other species.

## Acknowledgments

All movies and supplemental tables can be found on dropbox: https://goo.gl/GX4Szj. We thank the Duke University Genome Sequencing & Analysis Core for sequencing the RNA libraries and the Duke Light Microscopy Facility and Robertas Ursache for assistance in imaging. We thank Idan Efroni for assistance with the ICI pipeline, Guy Wachsmann for assistance in mapping RNAseq reads, Veronica G. Doblas for assistance with CIF2 treatments, Heather Belcher for technical assistance, Kim Gallagher for thoughtful discussion and members of the Benfey lab for comments on the manuscript. This research was funded by grants from the NIH (R01-GM043778) and the Howard Hughes Medical Institute and the Gordon and Betty Moore Foundation (GBMF3405) to PNB. N.G. was funded by an ERC Consolidator Grant (GA-N°: 616228 – ENDOFUN), an SNSF grant (31003A_156261). Additional support was provided to CD through an EMBO Short-term fellowship, to EES through startup funds from the University of Delaware, to PM by a FEBS Long-Term Fellowship, to TGA by an IEF Marie Curie fellowship

(Main Movies also available at https://goo.gl/GX4Szj)

**Movie 1** – CASP1 intensity fluctuates overtime. Max intensity projections of five day-old *pWER::*SHR-GFP roots (without CIF2) imaged overnight. Scale bars = 50µM.

**Movie 2** - CASP1 intensity depolarizes CASP1 channel in magenta. Sum intensity projections of five day-old *pWER::*SHR-GFP roots germinated on CIF2 then removed from CIF2 overnight. Scale bars = 50µM.

## Supplementary Materials

### List of Supplementary Materilas

**Figure S1**- *SHR*, *MYB36* but not *SCR*, can induce endodermal differentiation in the epidermis.

**Figure S2** - Epidermal expressed *SHR* and CIF2 treatment induce ectopic suberization.

**Figure S3** - *SGN3* is required but not sufficient for SHR and CIF2 induced endodermal reprogramming.

**Figure S4** - *MYB36* and *SCR* are not sufficient to induce complete endodermal reprogramming epidermis but *MYB36* and *SGN3* are required for SHR and CIF2-induced endodermal reprogramming.

**Figure S5** - Reprogrammed sub-epidermal layer resembles endodermis and retains some cortex gene expression.

**Figure S6** - Epidermal-expressed *SHR* can induce *SCR* and *MYB36*.

**Figure S7** - Ectopic suberization induced by *SHR* and CIF2 does not require SHR movement.

**Figure S8** - The reprogrammed sub-epidermal layer has limited barrier function.

**Figure S9** – *SHR* expressed in differentiated epidermal cells cannot induce CASP1

**Movie S1** – FDA rapidly penetrates sub-epidermal layer of *pWER::*SHR-GFP.

**Movie S2** - FDA rapidly penetrates sub-epidermal layer of *pWER::*SHR-GFP + CIF2.

**Table S1** – List of Differentially Expressed Genes among all RNAseq samples

**Table S2** – RNAseq Statistics

**Table S3** – Cloning Primers

**Supplementary References**

### Methods

#### Plant growth conditions

Seeds were gas sterilized, vernalized for 48h and plated on sterile ½X MS, 1% sucrose, 1% agar plates (for barrier assays and live imaging) or hydroponically as described in (*29*) (for clearing and staining), and grown under long-day conditions (16h light). Several plant growth conditions adjusting MS, sucrose, long day vs. 24h light, liquid and agar were tested and none alter the SHR- CIF2 induction phenotype. For transgenic lines containing the XVE inducible promoter system (*30*), seedlings were germinated on plates containing 10µM β-estradiol unless otherwise specified in captions. At four days post-stratification, seedlings were transferred to plates containing 100nM CIF2 (Invitrogen) or without CIF2 (including 10µM β-estradiol for XVE-containing seedlings), or CIF2 was added to liquid medium for 24h. For counting percentage of CASP positive cells in epidermis vs sub-epidermis, the 48h treatment was at three days post-stratification and 72h treatment at two days post stratification. All plants were imaged at five days old.

#### Clearing, staining and imaging of fixed samples

For lines in a CASP1-mCherry background, lignin was assessed by Basic Fuchsin staining as described in (*31*) and suberin was assessed by Fluorol Yellow staining as described in (*22*). Basic Fuchsin-stained roots were imaged on a Zeiss 510 confocal using a plan apochromatic 40x objective and Fluorol-Yellow stained roots on a Leica DM5000 microscope using a 10X objective (excitation/emission parameters below). For examining CASP1 expression, five day-old roots were fixed in PFA and cleared in Clearsee as described in (*32*). For lines in a *CASP1-GFP* background, lignin localization was assessed by combining Basic Fuchsin and ClearSee (*33*)). Suberin was assessed by combining Nile Red and ClearSee staining (*33*). Roots were imaged on a Zeiss 880 using a 40X or 20X objective. The following are the excitation and detection parameters: calcofluor white ex 405 nm, em 425–475 nm; ex Basic Fuchsin/mCherry 561 nm, em 600–650 nm; ex Fluorol-Yellow/GFP 488nm, em 525–550 nm; ex Nile Red 561 nm, em 600–620 nm.

#### Live-Imaging of CASP1 dynamics without CIF2 and after CIF2 treatment

Seeds were sterilized, stratified for 48h and plated on plates containing ½X MS, 1% sucrose, 1%agar with or without 100nM CIF2. At five days old, plants were removed from CIF2 and imaged on a Zeiss 880 for 16–24h on a ½X MS, 1% sucrose, 1% agar pad.

#### FDA and Propidium Iodide Assays

Seeds were sterilized, stratified for 48h and plated on sterile ½X MS, 1% sucrose plates, with or without 100nM CIF2 and with 10µM β-estradiol for seed lines containing the XVE-inducible promoter system. At five days old, a propidium iodide assay (*23*) or a FDA assay (*22*) was conducted.

#### FACS Sorting and RNA-sequencing

Seeds were sterilized with 3% (vol/vol) sodium hypochlorite and 0.1% Tween for 5 min and rinsed five times in sterile water. Seeds were stratified for 48h and plated on sterile 1X MS, 1% sucrose plates with or without 100nM CIF2. Roots were harvested on day 5, digested and subjected to Fluorescence Activated Cell Sorting (FACS) as described previously (*34*). Total RNA was isolated using RNeasy Micro-kit (Qiagen). RNA integrity and quantity were assessed on Agilent Bioanalyzer system and QuBit, respectively (ThermoFisher Scientific). cDNA synthesis for library preparation was carried out using the SMART-Seq v4 Ultra Low Input RNA kit for Sequencing (Takara). The Low Input Library Prep Kit v2 (Takara) was used to prepare libraries for sequencing. Single-end reads were obtained using the Illumina HiSeq4000 platform at the Duke University Sequencing Core.

#### Sequencing Analysis

All scripts for analysis can be found on github: https://github.com/cdrapek/Endodermis. Briefly, reads were aligned with TopHat and counts were normalized in EdgeR (*35*). Pearson correlation among libraries and mapping statistic can be found in Table S2. Differentially expressed gene lists (Table S1) were filtered by log-fold change with an absolutely value greater than or equal to 2, p-value of 0.01 and FDR of 0.01, aside from comparing *pWER::SHR-GFP* with and without CIF, in which the cut off was based solely on p-value. PCA analysis was carried out on the top 3000 genes with the most variation. Marker Spec scores and ICI values were calculated as previously described (*20*). Briefly, Spec scores were calculated from RootMap FPKM RNAseq data with the “getAllSpec” function and the default filters (medianfilter=0, cuts=FALSE, distshape=0). The index of cell identity (ICI) was calculated for each replicate using the “getIdentity” function. Values reported are for the normalized ICI score. Significance was determined from FDR corrected p-values.

#### Transgenic lines and cloning

The following lines have been previously published: p*WER*::SHR-GFP, CASP1-mCherry, CASP1-GFP, SGN3:venus, SCR: CFP. For full list of primers used for cloning, see supplemental material Table S3. The 3Kb fragment upstream of the *MYB36* start codon was fused to H2B and 3xYFP fragments using the multi-site gateway system. The inducible *WER-XVE* promoter was previously generated (*30*). The was fused to *MYB36* or *SCR* cDNA entry vectors and fluorophore-terminator containing entry vectors using the multi-site gateway system into a Norf-resistant destination vector. The *SHR-mKate2* lines were generated by fusing *SHR* cDNA to *mKate2* by In-Fusion technology (Takara) followed by insertion into pDONR221 vector by BP reaction. The p*WER-XVE* promoter and *SHR-mKate2* construct were assembly into the FastRed selection system destination vector using the Gateway Cloning system. For each transgenic line, four to eight T1 lines containing a single-insertion were observed and a representative line was selected for follow up study.

**Fig. S1.**
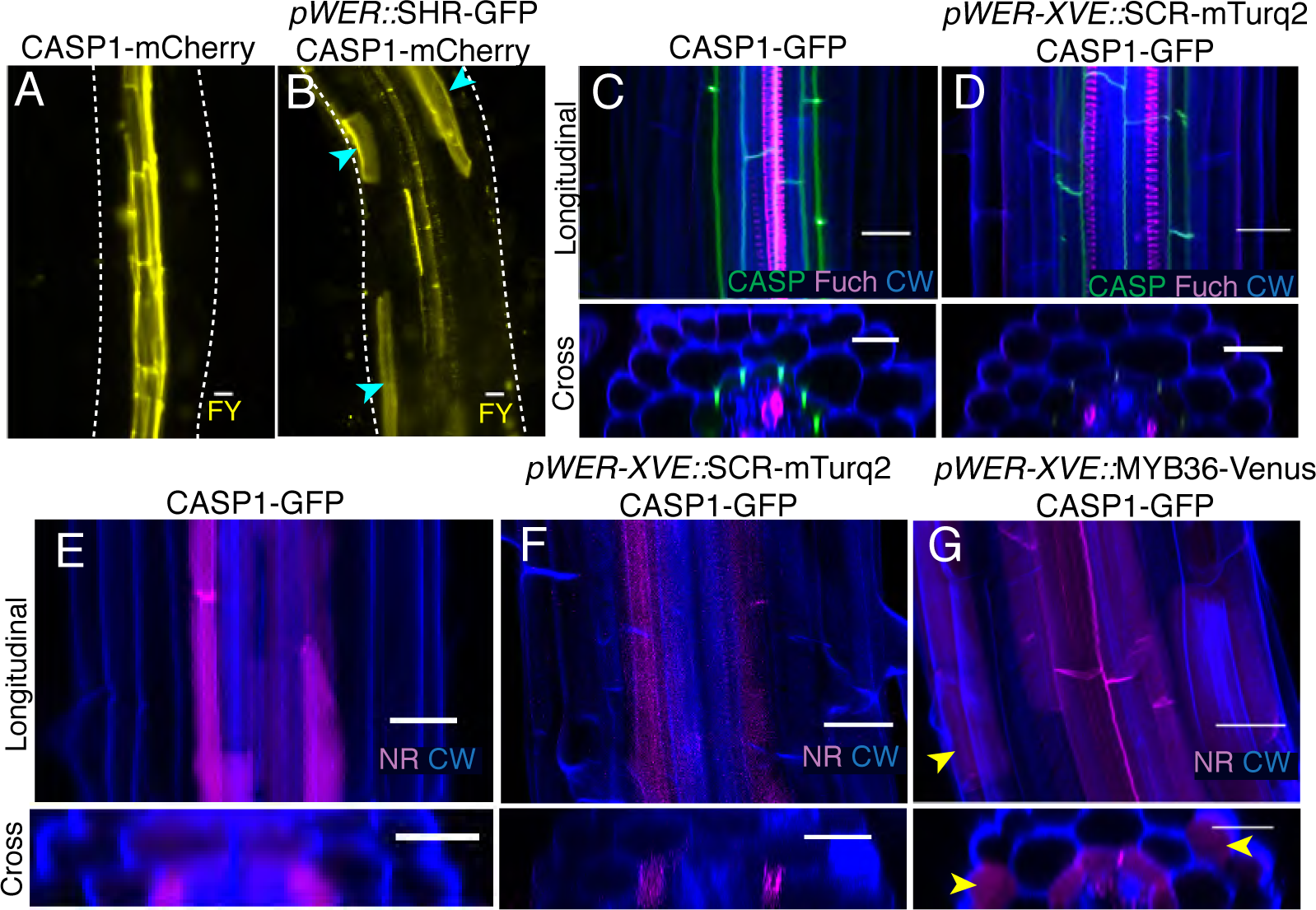
*SHR*, *MYB36* but not *SCR*, can induce endodermal differentiation in the epidermis. (A) Fluorol yellow (FY) staining of CASP1-mCherry (wild-type) roots. (B) Fluorol yellow (FY) staining of *pWER::*SHR-GFP x CASP1-mCherry roots. Cyan arrowheads indicate ectopic suberin. White dashed outlines in (A-B) outline root. (C) Sum projections of longitudinal and cross sections views of cleared CASP1-GFP roots germinated on 10µM β-estradiol stained with fuchsin (Fuch) and calcofluor white (CW). (B) Sum projections of longitudinal and cross sections views of cleared *pWER-XVE::SCR*-mTurquoise2 x CASP1-GFP roots germinated on 10µM β-estradiol stained with fuchsin (Fuch) and calcofluor white (CW). (E) Max projections of longitudinal and cross section views of cleared CASP1-GFP roots germinated on 10µM β-estradiol stained with Nile Red (NR) and calcofluor white (CW). (F) Max projections of longitudinal and cross section views of cleared *pWER-XVE::*SCR-mTurquoise2 x CASP1-GFP roots germinated on 10µM β-estradiol stained with Nile Red (NR) and calcofluor white (CW). (G) Max projections of longitudinal and cross section views of cleared *pWER-XVE::*MYB36-Venus x CASP1-GFP roots germinated on 10µM β-estradiol stained with Nile Red (NR) and calcofluor white(CW). Yellow arrowheads indicate cells with ectopic suberin in the epidermis. Scale bars = 20µM.

**Fig. S2.**
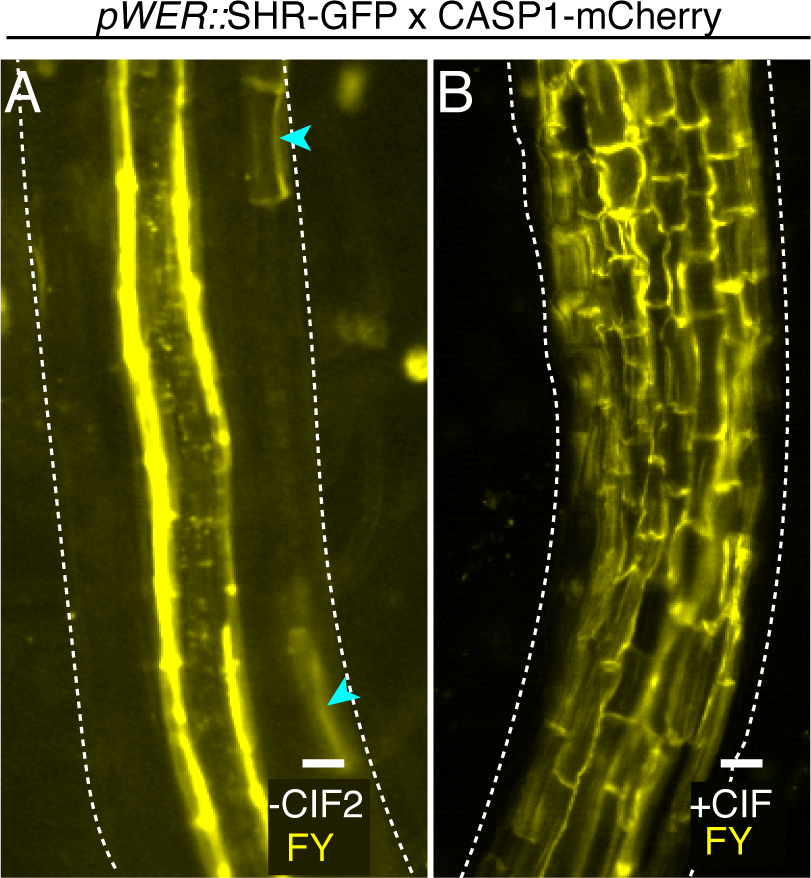
Epidermal expressed *SHR* and CIF2 treatment induce ectopic suberization. (A) Longitudinal views of fluorol yellow staining (FY) of *pWER::*SHR-GFP without CIF2. Cyan arrowheads indicate cells with ectopic suberin deposition. (B) Longitudinal views of fluorol yellow staining (FY) *pWER::*SHR-GFP treated with CIF2. White dashed lines indicate outline of the root. Scale bars = 20µM.

**Fig. S3.**
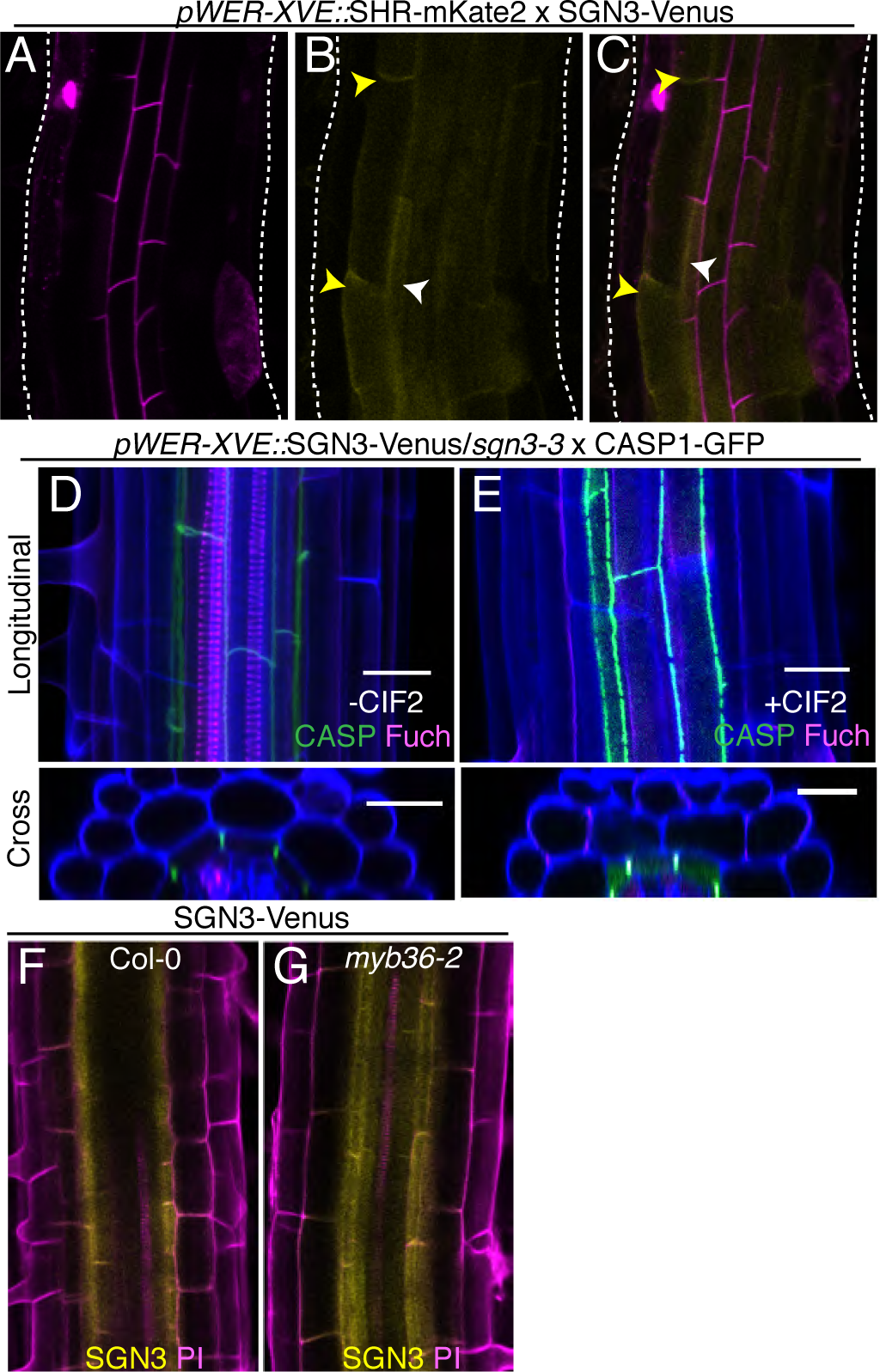
*SGN3* is required but not sufficient for SHR and CIF2 induced endodermal reprogramming. (A) Sum projections of live imaged *pWER-XVE::*SHR-mKate2 x SGN3-Venus roots germinated on 10µM β-estradiol. Yellow arrowheads indicate ectopic SGN3, white arrowheads indicate endogenous SGN3. (D) Sum projections of longitudinal and cross sections views of cleared *pWER-XVE::*SGN3-Venus/*sgn3-3* x CASP1-GFP roots germinated on 10µM β-estradiol without CIF2 and with 100 Nm CIF2 (E), stained with fuchsin (Fuch) and calcofluor white (CW). (F) Expression of SGN3-Venus in wild-type Propidium Iodide (PI) stained roots and (G) *myb36-2* roots.

**Fig. S4.**
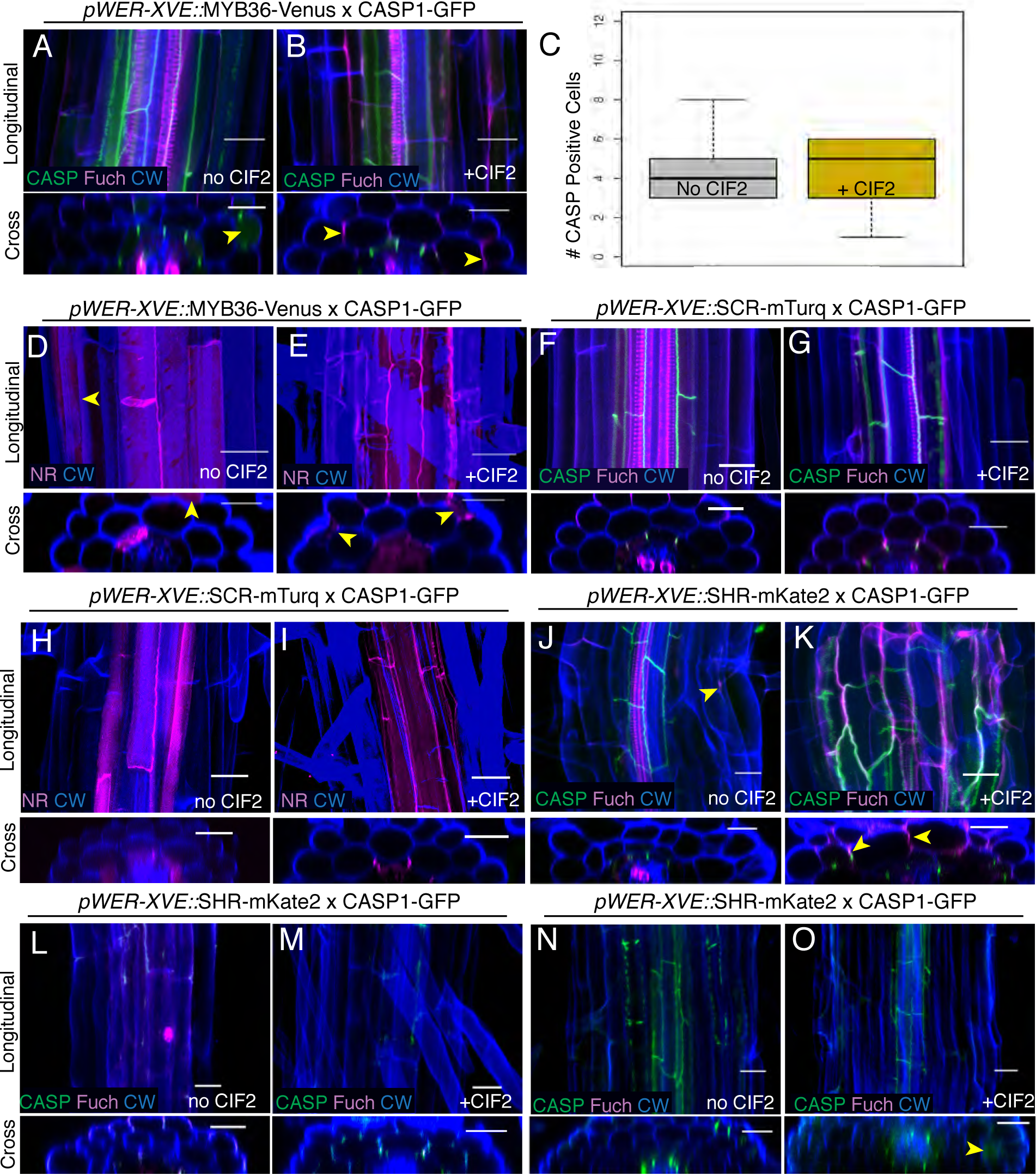
*MYB36* and *SCR* are not sufficient to induce complete endodermal reprogramming but *MYB36* and *SGN3* are required for SHR and CIF2-induced endodermal reprogramming. (A) Sum projections of longitudinal and cross section views of cleared *pWER-XVE::*MYB36-Venus x CASP1-GFP roots germinated on 10µM β-estradiol without CIF2, or (B) with 100nM CIF2, stained with fuchsin (Fuch) and calcofluor white(CW). Yellow arrowheads indicate cells with ectopic CASP1 or lignin. (C) Quantification of the number of cells expressing ectopic CASP1 in *pWER-XVE::*MYB36-Venus x CASP1-GFP roots germinated on 10µM β-estradiol with or without out CIF2. Standard t-test p-value = 0.77. (D) Sum projections of longitudinal and cross section views of cleared *pWER-XVE::*MYB36-Venus x CASP1-GFP roots germinated on 10µM β-estradiol without CIF2, or (E) with 100 nM CIF2, stained with Nile Red (NR) and calcofluor white(CW). Yellow arrowheads indicate cells with ectopic suberin. (F) Sum projections of longitudinal and cross section views of cleared *pWER-XVE::*SCR-mTurquoise roots x CASP1-GFP germinated on 10µM β-estradiol without CIF2, or (G) with 100 nM CIF2, stained with fuchsin (Fuch) and calcofluor white(CW). (H) Sum projections of longitudinal and cross section views of cleared *pWER-XVE::*SCR-mTurquoise roots x CASP1-GFP germinated on 10uM β-estradiol without CIF2, or (I) with 100nM CIF2, stained with Nile Red (NR) and calcofluor white(CW). (J) Sum projections of longitudinal and cross section views of cleared *pWER-XVE::*SHR-mKate2 x CASP1-GFP roots germinated on 10µM β-estradiol without CIF2, or (K) with 100 nM CIF2, stained with fuchsin (Fuch) and calcofluor white(CW). Yellow arrowheads indicate examples of ectopic CASP1 or lignin. (L) Sum projections of longitudinal and cross section views of cleared *pWER-XVE::*SHR-mKate2/*myb36-2* x CASP1-GFP roots germinated on 10µM β-estradiol without CIF2, or (M) with 100 nM CIF2, stained with fuchsin (Fuch) and calcofluor white(CW). (N) Sum projections of longitudinal and cross section views of cleared *pWER::*SHR:GFP/*sgn3-3* x CASP1-GFP germinated on 10µM β-estradiol without CIF2, or (O) with 100 nM CIF2, stained with fuchsin (Fuch) and calcofluor white(CW). Scale bars = 20µM.

**Fig. S5.**
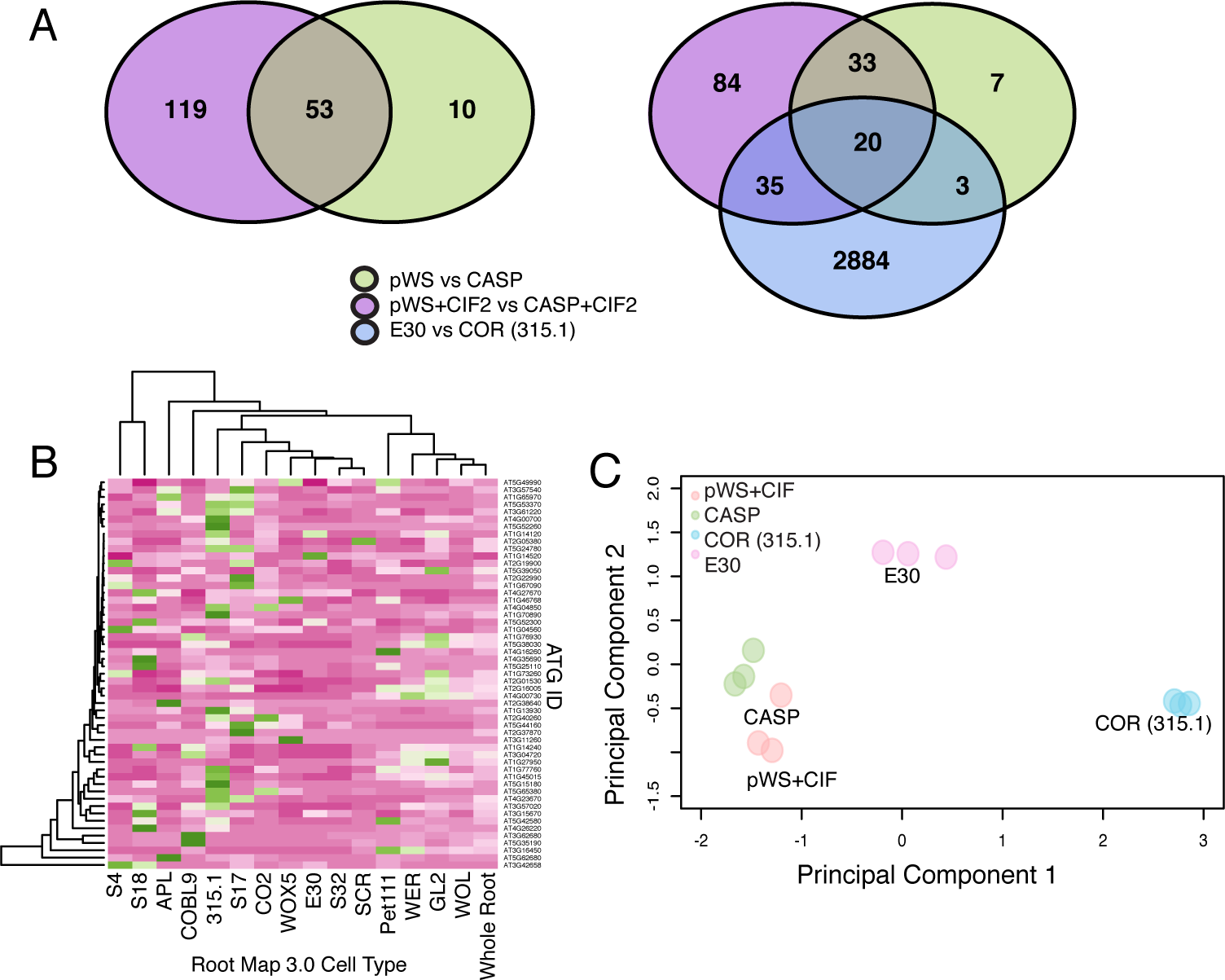
Reprogrammed sub-epidermal layer resembles endodermis and retains some cortex gene expression. (A) Venn diagram of overlap of differentially expressed genes between samples in Fig 2(E). (B) Heat map of native the 127 genes differentially expressed between *pWER::*SHR-GFP samples and wild-type samples. (C) Principal component analysis of *pWER::*SHR-GFP samples treated with CIF2 compared to wild-type CASP, E30 and COR(315.1) samples.

**Fig. S6.**
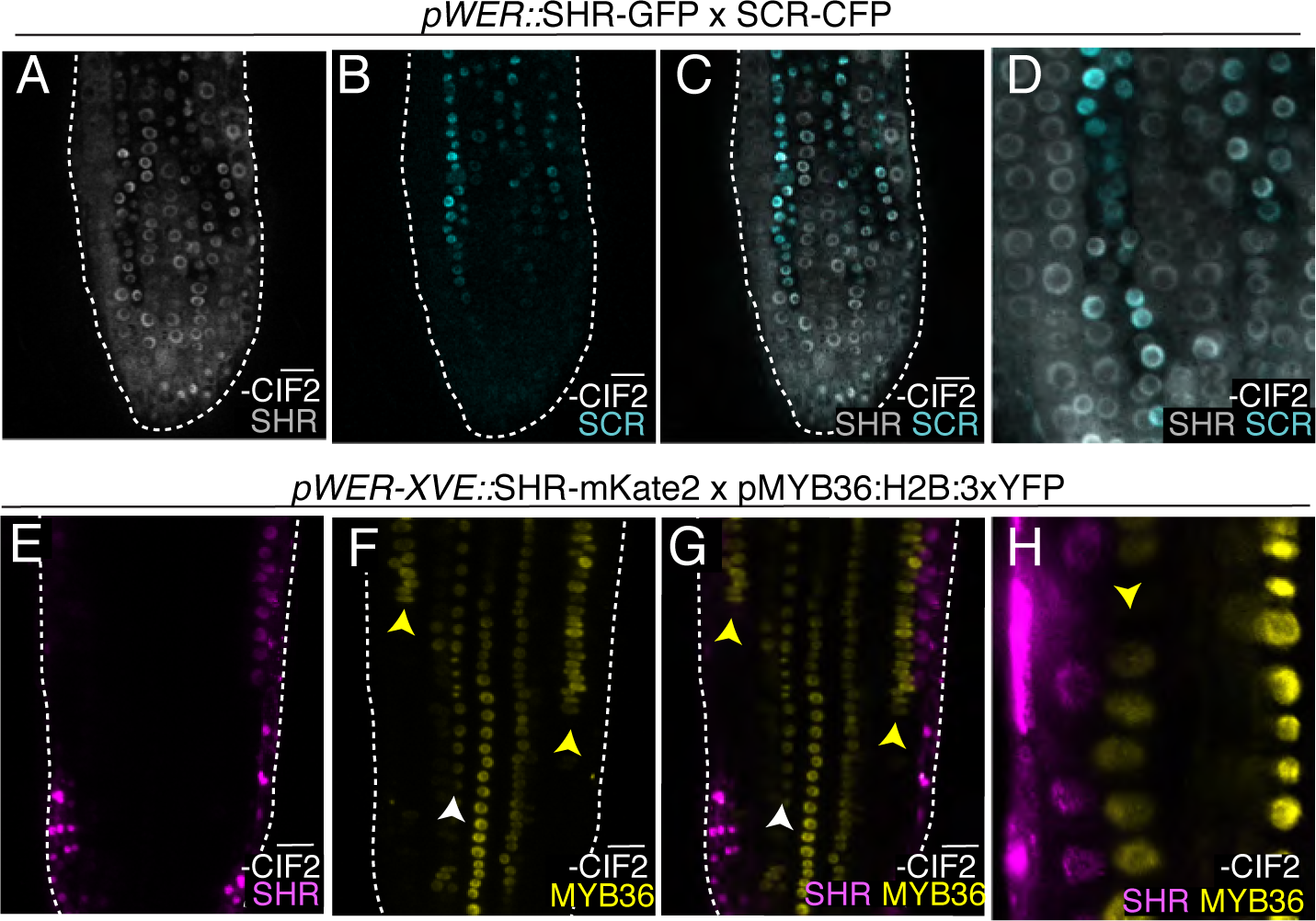
Epidermal-expressed *SHR* can induce *SCR* and *MYB36*. Sum projection of meristematic epidermal cells of *pWER::*SHR:GFP x SCR-CFP roots (A) GFP (SHR) channel, (B) CFP (SCR) channel, (C) merge, (D) zoomed in. Sum projections of meristematic epidermal cells of *pWER-XVE::*SHR-mKate2 x pMYB36:H2B:3XYFP roots germinated on 10µM β-estradiol (E) mKate2 (SHR) channel, (F) YFP (MYB36) channel, (G) merge, (H) zoomed in. Scale bars = 20µM. Yellow arrowheads indicate examples of ectopic expression of *SCR* or *MYB36*. White arrowheads indicate examples of endogenous expression of *SCR* or *MYB36.* White dashed lines represent root outline.

**Fig. S7.**
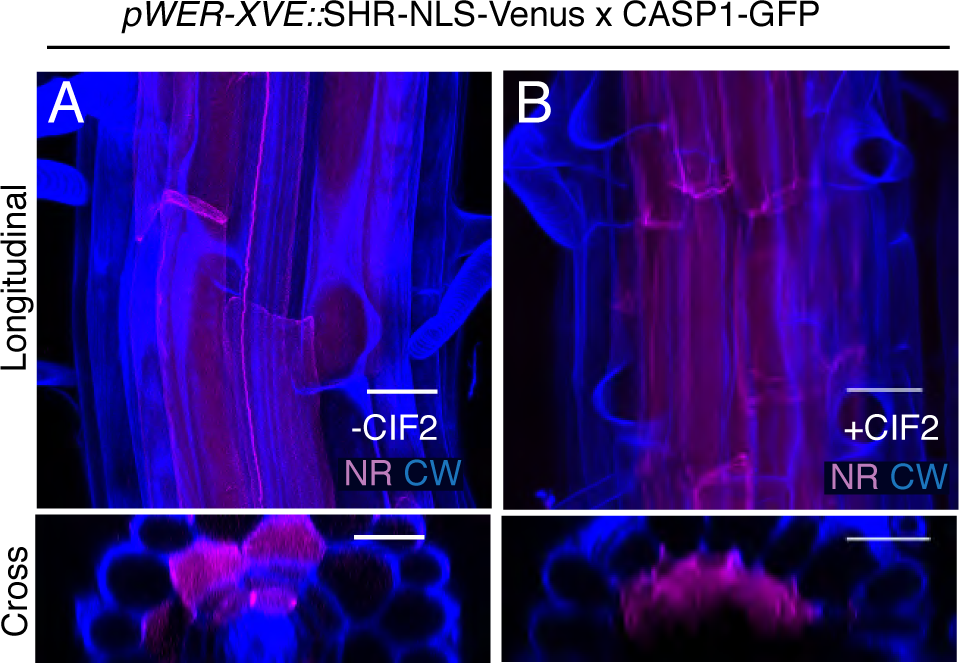
Ectopic suberization induced by *SHR* and CIF2 does not require SHR movement. (A) Max projections of longitudinal and cross section views of cleared *pWER-XVE::*SHR-NLS-Venus roots germinated on 10µM β-estradiol cleared and stained with Nile Red (NR) and calcofluor white (CW). (B) Max projections of longitudinal and cross section views of cleared *pWER-XVE::*SHR-NLS-Venus germinated on 10µM β-estradiol and 100 nM CIF2 cleared and stained with Nile Red (NR) and calcofluor white (CW). Scale bars = 20µM.

**Fig. S8.**
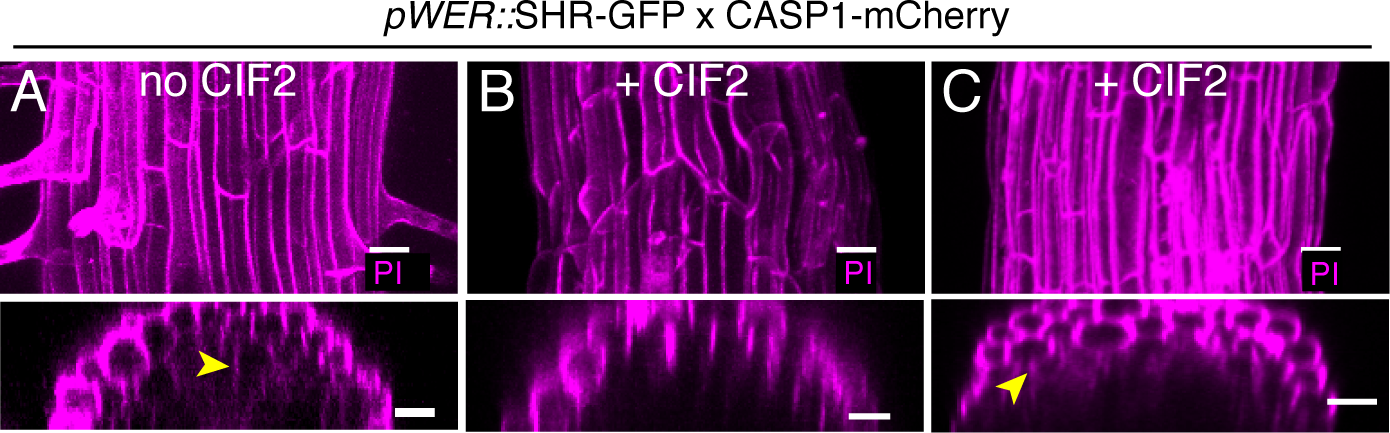
The reprogrammed sub-epidermal layer has limited barrier function. (A) Max projections of longitudinal and cross section views of *pWER::*SHR-GFP roots (no CIF2) stained with Propidium iodide. Yellow arrowhead indicates example cell stained with PI below sub-epidermal layer. (B) Max projections of longitudinal and cross section views of a *pWER::*SHR-GFP root treated with CIF2. (C) An additional example of max projections of longitudinal and cross section views of a *pWER::*SHR-GFP root treated with CIF2. Yellow arrowhead indicates example cell stained with PI below sub-epidermal layer. Scale bars = 20µM.

**Fig. S9.**
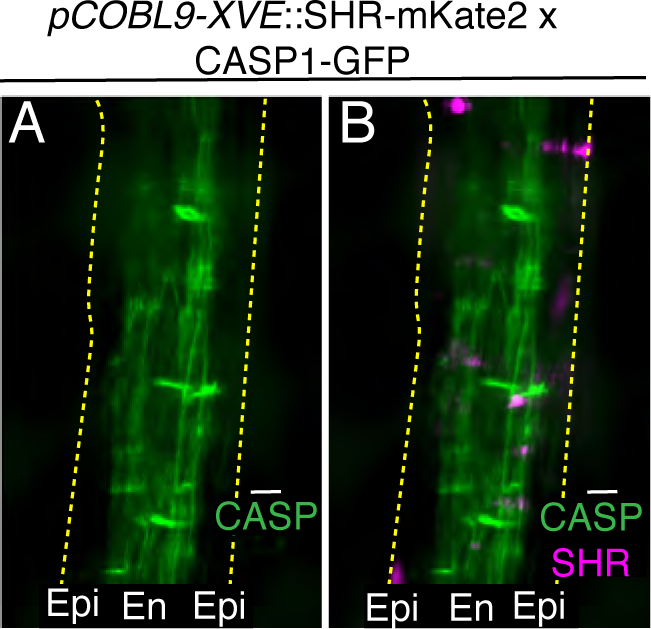
*SHR* expressed in differentiated epidermal cells cannot induce CASP1. Dissecting microscope images of *pCOBL9-XVE::*SHR-mKate2 T1 example treated for 2 days on 10µM β-estradiol (A) CASP1-GFP channel and (B) SHR-mKate2, CASP1-GFP merge. Epi = epidermis, En = endodermis, yellow dashed lines represent outline root. Scale bars = 50µM.

(Supplemental Movies available at https://goo.gl/GX4Szj)

**Movie S1** – FDA rapidly penetrates sub-epidermal layer of *pWER::*SHR-GFP. Sum projections of live imaging of FDA application to *pWER::*SHR-GFP roots treated without CIF2 treatment. Scale bars =50µM.

**Movie S2** - FDA rapidly penetrates sub-epidermal layer of *pWER::*SHR-GFP + CIF2. Sum projections of live imaging of FDA application to *pWER::*SHR-GFP roots treated with CIF2. Scale bars = 50µM.

**Supplemental Tables** available at https://goo.gl/GX4Szj

## Author contributions

CD, PNB and NG designed experiments. CD and PNB wrote the manuscript. CD carried out CIF2 treatments, barrier assays, FDA assays, staining/clearing and cell sorting. CD and EES carried out RNA-seq analyses. CD and PM carried out live imaging experiments. All authors contributed to discussion and interpretation of results. CD, TGA and JHH generated the transgenic lines.

